# Mechanical scarification, seed origin, and substrate influence on germination of *Samanea tubulosa*

**DOI:** 10.1101/2020.02.10.940759

**Authors:** Leonardo L. P. Regnier

## Abstract

*Samanea tubulosa* is an important species mainly to its uses in reforestation projects and as a wood and fruit resource. This species seed dormancy and other scarce germination information have been limiting the adequate seedling production process. Thus, this study tried to understand how seed origin, mechanical scarification, and substrate could influence the seedling production process. The population affected the time required for germination and germination speed, probably due to high genetic variability. Whereas mechanical scarification did not present statistical differences in the control treatment, with seeds in the natural condition, indicating that seed dormancy could be variable. Substrate promoted significant differences in germination rate. The population seems to affect, but not preclude seedling production. Mechanical scarification did not provide a relevant enhancement of germination. Vermiculite presented a significantly higher germination rate than the organic-based substrate.

## 1. Introduction

Samanea tubulosa (Benth.) Barneby & J. W. Grimes, popularly known in Brazil as *Sete-cascas*, and *Saman* in Spanish America, is an arboreous and deciduous tree. This species is native to South America and presents high tolerance to sunlight and the natural occurrence fires of Cerrado, the Brazilian savannah. It is characterized as a pioneer species, colonizing glades and open fields [1]. This species is commonly used in urban afforestation and reforestation programs, especially in disturbed areas [2]–[5].

Fruits are the main source of this species. They are usually employed as cattle food and when empowered are applied to goat and chicken feeding because of their high protein and sugar content [1]. Fruits can be eaten by humans and are usually utilized in cachaça production, resulting in *“Aguardente-de-saman”*. The wood from this species is widely exploited, especially in the furniture industry and fences [6]. Alcohol production, timber, and paper production are other recent uses of this species [1].

Seed dormancy is characterized by seeds that even when exposed to favorable environmental conditions, such as light exposure, moisture, oxygen, and temperature, they do not germinate [7], [8]. Seed dormancy is a common problem in the seedling production of arborous trees, and especially frequent in the Fabaceae family [7], [9]–[11], being nocuous to the seedling production process because fast and uniform germinations are required [10].

Understanding seed behavior from seeding until seedling establishment is also important for the successful application of wild species in reforestation programs [12], [13]. Thus, recognizing techniques that could improve the seedling production process is important [10],[11]. Especially of *S. tubulosa* since it is one of the most indicated species to use in this kind of project and urban afforestation.

The *S. tubulosa* high-level dormancy is already recognized as a result of its impermeable tegument [1], [6], [7]. In general, in the natural environment, temperature variations, acid soil, microorganism metabolism, and animal consumption of fruits are the main methods to overcome dormancy [11]. Chemical scarification using acid compounds is the most common pre-germinative procedure to overcome *S. tubulosa* seed dormancy. However, this seed dormancy overcoming procedure is very expensive and imposes some practical obstacles. In general, these compounds are very difficult to manage when a large number of seeds are submitted to this process, present the risk of accidents [9], [11], or even generate wounds to seed embryonic tissues [7], promoting abnormalities in seedlings [14]. Alternative methods, such as hot water treatment, are mentioned as not adequate to overcome *S. tubulosa* seed dormancy [10], [11].

Mechanical scarification is recommended by some authors [4], [7], [11], [15] as a viable alternative to chemical scarification. However, these authors recommend manual scarification techniques with sandpaper and discourage the use of the rotary drum because of the great damage that this last technique could provide to the embryo. Oliveira et al. [10] also measured the impact of mechanical scarification of *S. tubulosa* with scissors. In addition, these mentioned mechanical methods are infeasible in the seedling production context, which requires the processing of an extensive number of seeds without taking too much time.

In general, to forestry seeds, the parent plants seem to influence seedling quality [16]–[18]. Affecting some germination parameters, such as the total germination rate [19], the time required for germination, or even both [18]. This aspect is very important to be analyzed because, in the seedling production context, the choice of a progenitor population used as seed sources could affect seedling production [16], [17].

The substrate is essential for seedling germination [11]. Conflicting information could also be found about the best germination substrate, while some authors point out that to *S. tubulosa* better germination indexes were obtained when sowed in the organic-based substrate [20], others affirm that vermiculite is the best substrate for *S. tubulosa* production [11]. However, proper comparison between germination on inert substrates such as sand or vermiculite and in the organic-based substrate has not been extensively explored.

Thus, this study focused on understanding how mechanical scarification, seed origin, and substrate could influence *Samanea tubulosa* germination.

## 2. MATERIAL & METHODS

This study was conducted at the Harry Blossfeld plant nursery of São Paulo, situated in the city of Cotia (23°36’30.0”S 46°50’48.9”W). According to Köppen’s climate classification, the study area presents Cwa, an altitude tropical climate [21]. Featuring moderate and concentrated rains during summer, dry winter [21], and the highest mean temperature above 22°C.

Plant material was collected in the city of São Paulo between September and October of 2018 from two different populations about 125 km away from each other. The harvesting method consisted of mature fruits gathered directly from the ground. The material was kept in open plastic bags at room temperature for 15 days to recognize the emergence of insects and/or pathogens [17]. After this period, fruits were opened and seeds were removed, as indicated by Carvalho [1], using a rubber hammer.

Population influence on germination was measured using 630 seeds from population I and 126 from population II. These seeds, without treatment, were planted in 280 cm^3^ greenhouse tube, with an organic-based substrate composed by 2 liters of local horizon B soil, 8 liters of rice husk, 5 liters of Basaplant^®^ forestry substrate, 46.5g of Yoorin K^®^ potassium thermosetting fertilizer, and 46.5 g of gradual release fertilizer Osmocote Plus^®^.

Evaluation of the seed processing method involved a sample of 252 scarified seeds and 630 seeds in a natural condition, planted in the previously mentioned organic-base substrate and greenhouse tubes. The mechanical scarification procedure consisted of the use of electronic emery until conspicuous seed tegument rupture of the opposite side of the mycropile.

Seeking to test substrate influence, 252 scarified seeds were sowed in the same previously mentioned organic-based substrate and the greenhouse tubes. While 1452 scarified seeds were seeded in white polypropylene trays containing vermiculite as the substrate.

After all the mentioned treatments, the material was kept in a greenhouse with white plastic covering and a fogging watering system with periodic activation every 35 min. Plant emergence was recorded about 20 days after seeding, with measurements nearly every 7 days.

All data were compiled into the Excel component of the Microsoft Corporation Office pack. The most common germination indexes were obtained using GerminaQuant software [22]. All statistical tests were executed with the data of the repetitions, also using the GerminaQuant software, with ANOVA followed by the Tukey test adopting a critical p-value of 5% (α < 0.05).

## 3. RESULTS AND DISCUSSION

### Population origin

The population seems to influence the time required for *S. tubulosa* seed germination (Figure 1) since we observed only differences between the mean germination rate (Figure 1B), mean germination time (Figure 1C), and Germination speed (Figure 1D). Germination rate (Figure 1A) was not affected by seed origin during the analyzed period. These same indexes were also affected when comparing the population influence to *Pterocarpus rohrii*, another species of the Fabaceae family [18].

**Figure 1.**
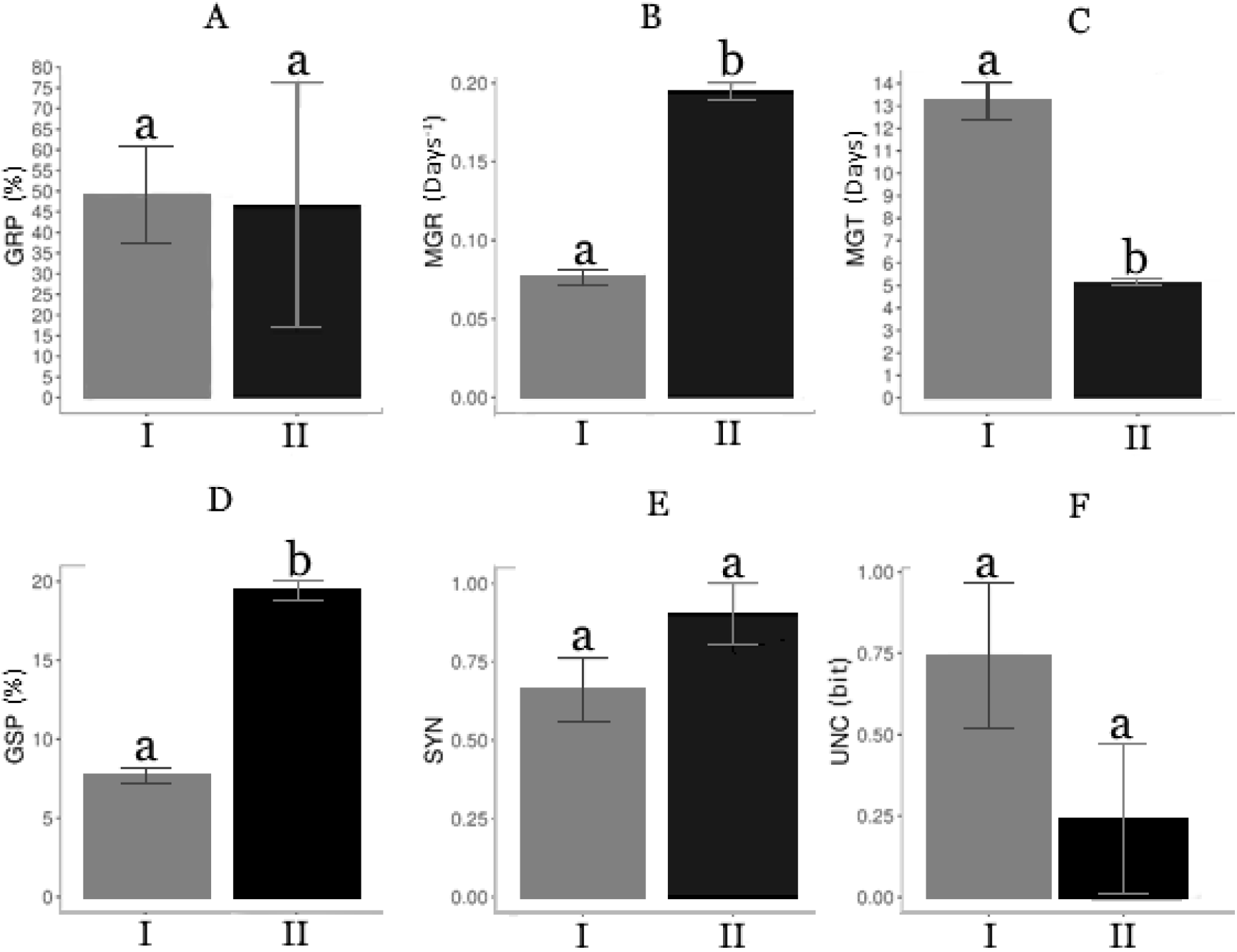
**A-** Germination Proportion (GRP), **B-** Mean germination rate (MGR), **C-** Mean germination time (MGT), **D-** Germination speed (GSP), **E-** Synchronization index (SYN) and **F-** Uncertainty index (UNC), according to seed origin: population I (I), population II (II). Lowercase letters present statistical differences of p<0.05. Error bars present the standard error mean.

Mean germination time (MGT) represents the mean time required to germinate one seed of a seed lot. In this study, population II presented the double value of germination speed (GSP), and its seeds take about 5 days to germinate. Whereas in population I, the mean time required for one seed germinate (MGT) comprehends 13 days, almost three times the obtained value of population I. To seedling production process, differences of 7 days are relevant but not crucial [18]. Thus, the seedling production process could be affected by seed origin, but it seems that it is not substantially impaired to this influence. Similar results were found for *Hymenaea courbaril* [23], but opposite findings to *P. rohrii* [18], both members of the same family.

In general, variations between populations are expected to non-domesticated species [17]. Wild species present greater regional variations because they are under a continuous evolutionary process, and this could create great variations between populations [17], [18], [24]. Therefore, depending on the study sampling standardization, the results will be only regionally applicable [17].

Our results indicate that, to *S. tubulosa*, choosing a progenitor population seems to affect the time indexes, but not the germination proportion, faster germinations are better, reducing the time a seed lot occupies in the greenhouse or the production area. Thus, choosing an adequate parent population could provide a slight reduction in the required time for seed lot germination.

The main differences between the germination patterns were seedling emergence (Figure 2). Both populations reached the same germination amount at the end of this experiment. In addition, seeds from population I presented faster emergence (also indicated by the indexes). It is important to note that a greater time of experiment could provide different results. Shorter evaluations under 9 days could lead to the erroneous conclusion that population I seeds almost did not germinate. Therefore, as mentioned before, it is important to consider time during germination evaluations [18], [25].

**Figure 2.**
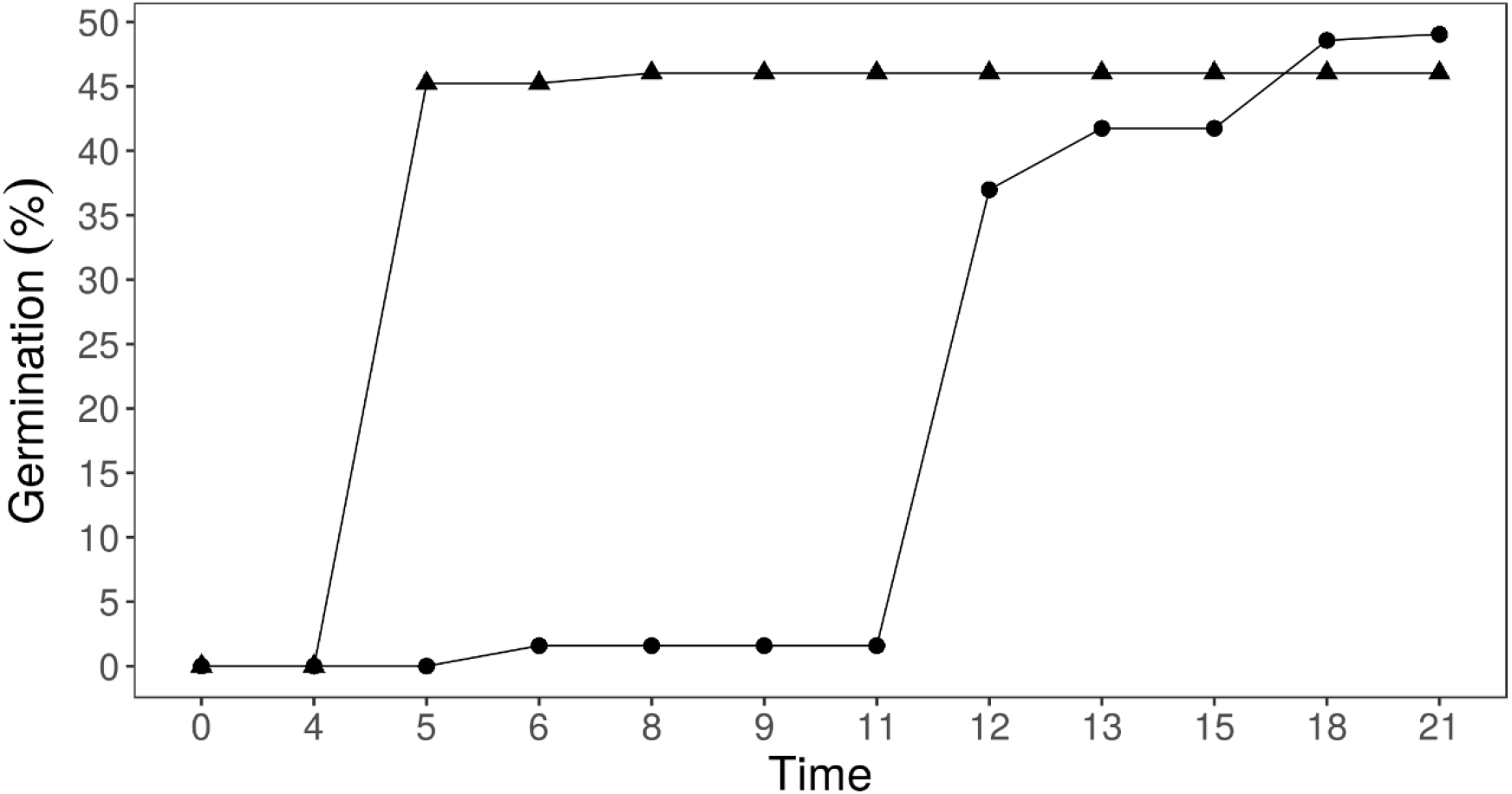
Cumulative germination of *S. tubulosa* according to seed origin: Population I (●), Population II (▲).

### Seed processing method

Mechanical scarification with emery and seeds in natural conditions did not present significant differences in any of the studied indexes (Figure 3). Variations in germination were greater in non-processed seeds in germination proportion (Figure 1A), synchrony (Figure 1E), and uncertainty (Figure 1F) indexes. Indicating that when seeds are not scarified, they present great variations between seed lots to those mentioned indexes. Great uniformity promoted by scarification has already been recognized. Especially because, during the imbibition phase, water is the main agent to promote germination, as scarification facilitates water uptake, higher values of uniformity are obtained [26].

**Figure 3.**
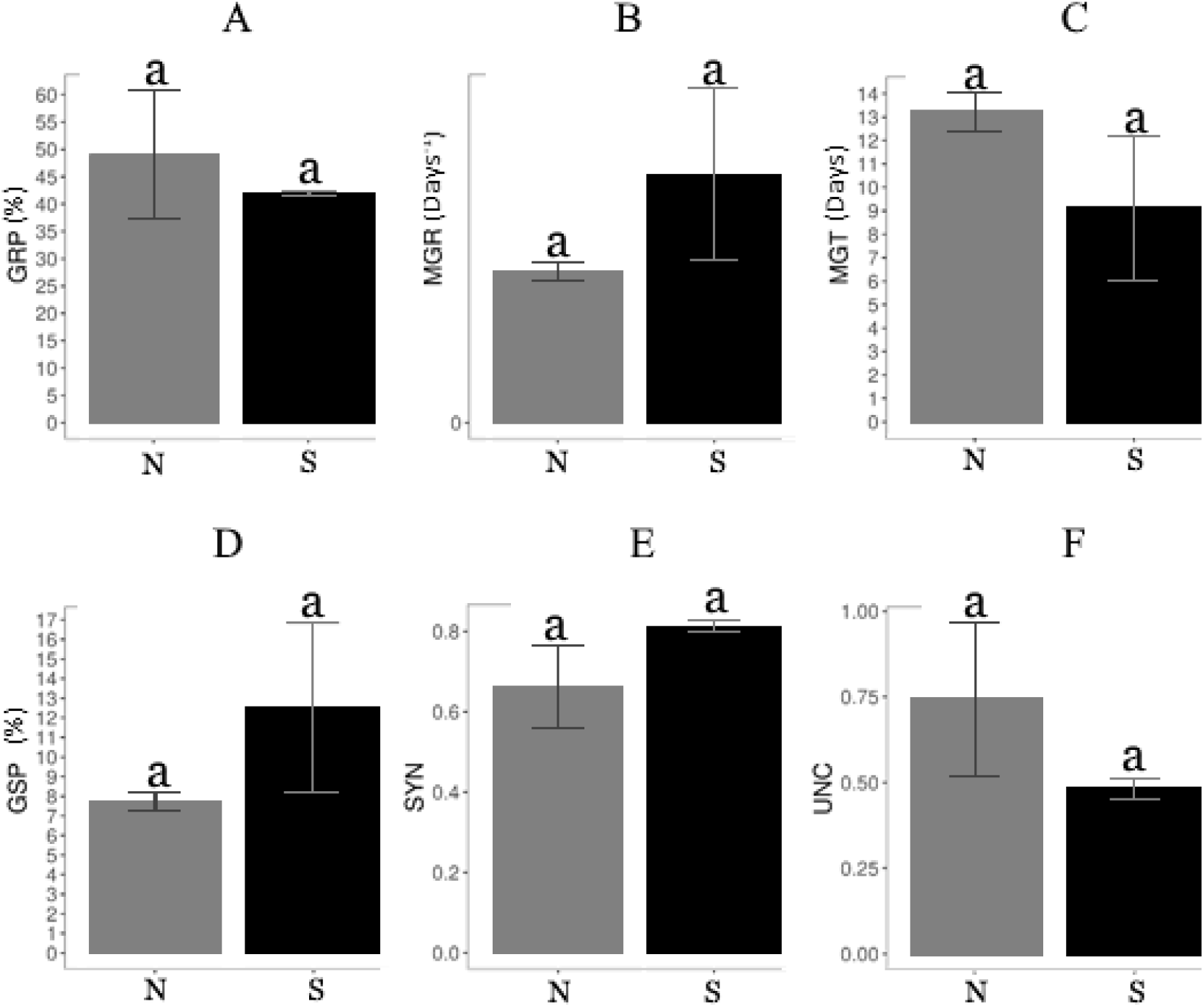
**A-** Germination Proportion (GRP), **B-** Mean germination rate (MGR), **C-** Mean germination time (MGT), **D-** Germination speed (GSP), **E-** Synchronization index (SYN), and **F-** Uncertainty index (UNC) according to seed processing method: Scarified (S), and non-processed seeds (N). Lowercase letters present statistical differences using p<0.05. Error bars present the standard error mean.

Other studies showed that *S. tubulosa* acid-scarified seeds presented germination rates higher than 85%, and seeds under natural condition reached 55% [4]. These values are very discrepant from the data we obtained for mechanically scarified seeds. In general, both scarified and natural condition seeds reached approximately 50% germination. Although these authors promoted mechanical scarification using sandpaper, while in this study we promoted scarification by emery. Martins et al. [9] mentioned that scarification with sandpaper efficiency is highly variable according to the person dexterity to execute the procedure. However, the genetic variations found in wild species, the evaluation time of two months, and the controlled temperature in the mentioned study [4] could not be rejected as possible reasons for the discrepancies. It is already known that temperatures higher than 25°C promote better germination rates in *S. tubulosa* [7]. Thus, as those authors adopted 30°C during the experiment, this could explain the greater results than the present study.

Other authors have already mentioned that there is a slight increase, not always significant, on germination speed when seeds are mechanically scarified [4], [13]. We found a similar pattern, although without statistical significant.

Seed dormancy and germination are also affected by the genetic diversity of native species [17], [18]. In general, in the Fabaceae family, the dormancy level could vary between populations [27] or even in the same seed lot [28]. This effect could be associated with tegument thickness and waterproofing compounds, resulting in different degrees of seed dormancy [10], [13]. Thus, beyond the influence of the analysis period mentioned before, variations between seed dormancy could also interfere with germination.

Seeds in the natural condition presented a delayed emergence when compared to the scarified seeds (Figure 4). However, at the end of this study analysis, the germination pattern of the treatments had close values of germination rates. This general pattern was also obtained by Cruz et al. [28] to *H. intermedia*, another member of the Fabaceae family, that present seeds with different dormancy degrees [27]. Mechanical scarification also did not promote enhanced germination of *S. tubulosa* in the study conducted by Barbosa et al. [13].

**Figure 4.**
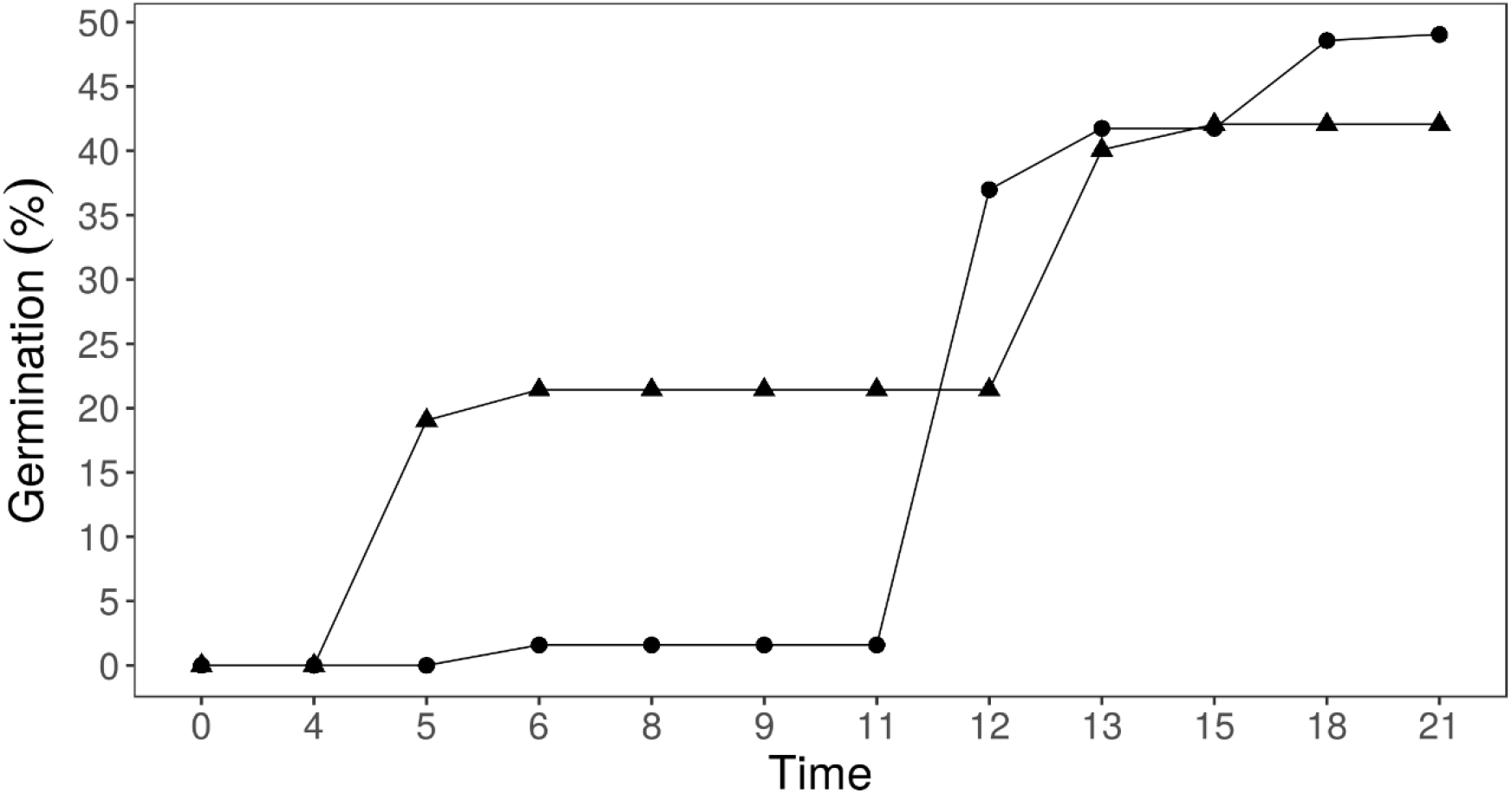
Cumulative germination of *S. tubulosa* according to seed processing method: seeds in natural conditions (●), and scarified seeds (▲).

Carvalho [1] indicates that scarified seeds’ emergence occurs between 14 and 42 days after sowing, reaching about 99% of germination. While non-processed seeds’ emergence occurred between 40 and 90 days obtaining 2% germination. In the present study, the results were very discrepant when compared to the data presented by this author. We found that scarified seeds presented the first emergencies on the fifth day, reaching approximately 49% of the germination rate (Figure 4). Whereas seeds in natural conditions presented the first emergencies on the sixth day after sowing, with 42% germination. Our results are closer to Oliveira et al. [10] data, that the first emergence of non-processed seeds is about the tenth day, and scarified seeds three days after seeding. In general, great variations between seed lots and different climatic regions could be responsible for the discrepancies in some experimental data.

### Germination substrate

Substrate seems to influence only the germination proportion (Figure 5A). All the other studied indexes were not consistently affected. Vermiculite seems to be a better option since it presented a greater germination rate than the organic-based substrate. First, Carvalho [1] found that *S. tubulosa* easily developed in the sand, indicating that inert substrates probably promote better germination. Sales [11] measured the differences between inert substrates of sand and vermiculite, and this last substrate presented the best results. Subsequently, Gonçalves [20] compared different compositions of organic-based substrates but did not measure the differences between vermiculite and his results. This present study emphasizes Carvalho [1] findings and the hypothesis that vermiculite is the best substrate for *S. tubulosa* germination, even when compared to the organic-based substrate.

**Figure 5.**
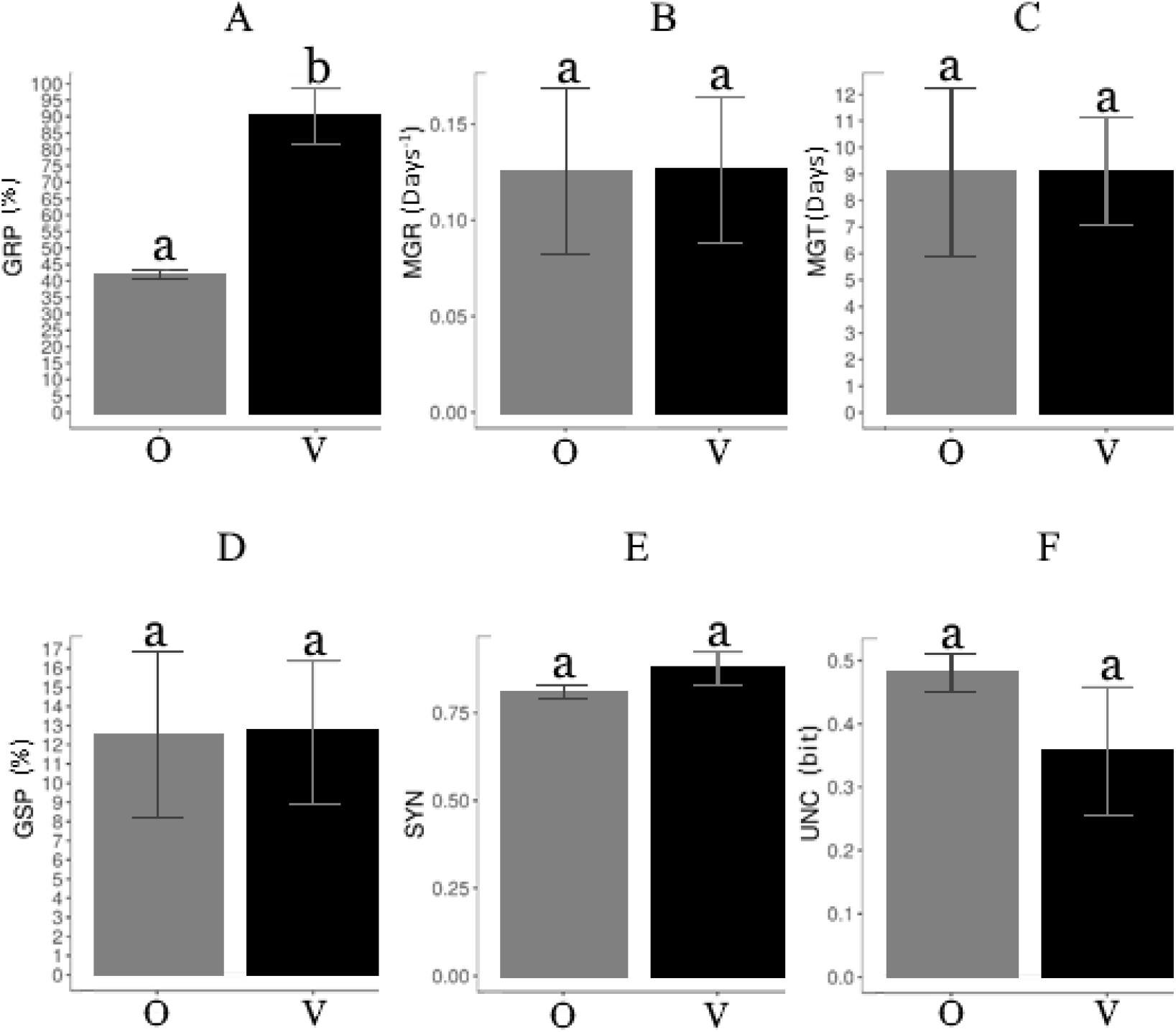
**A-** Germination Proportion (GRP), **B-** Mean germination rate (MGR), **C-** Mean germination time (MGT), **D-** Germination speed (GSP), **E-** Synchronization index (SYN), and **F-** Uncertainty index (UNC) according to the substrate: Organic-based (O), and vermiculite (V) substrate. Lowercase letters present statistical differences using p<0.05. Error bars present the standard error mean.

Germination pattern seems to accumulate differences over time, being clear the split promoted by those methods at the end of the analyzed period (Figure 6). The substrate can influence germination because aeration, structure, water retention, pathogenic infestation, and other aspects can be highly different [29], [30]. It is important to note that the cumulative germination curves were very similar in shape, but vermiculite promoted an upper germination proportion at the end of the study analysis.

**Figure 6.**
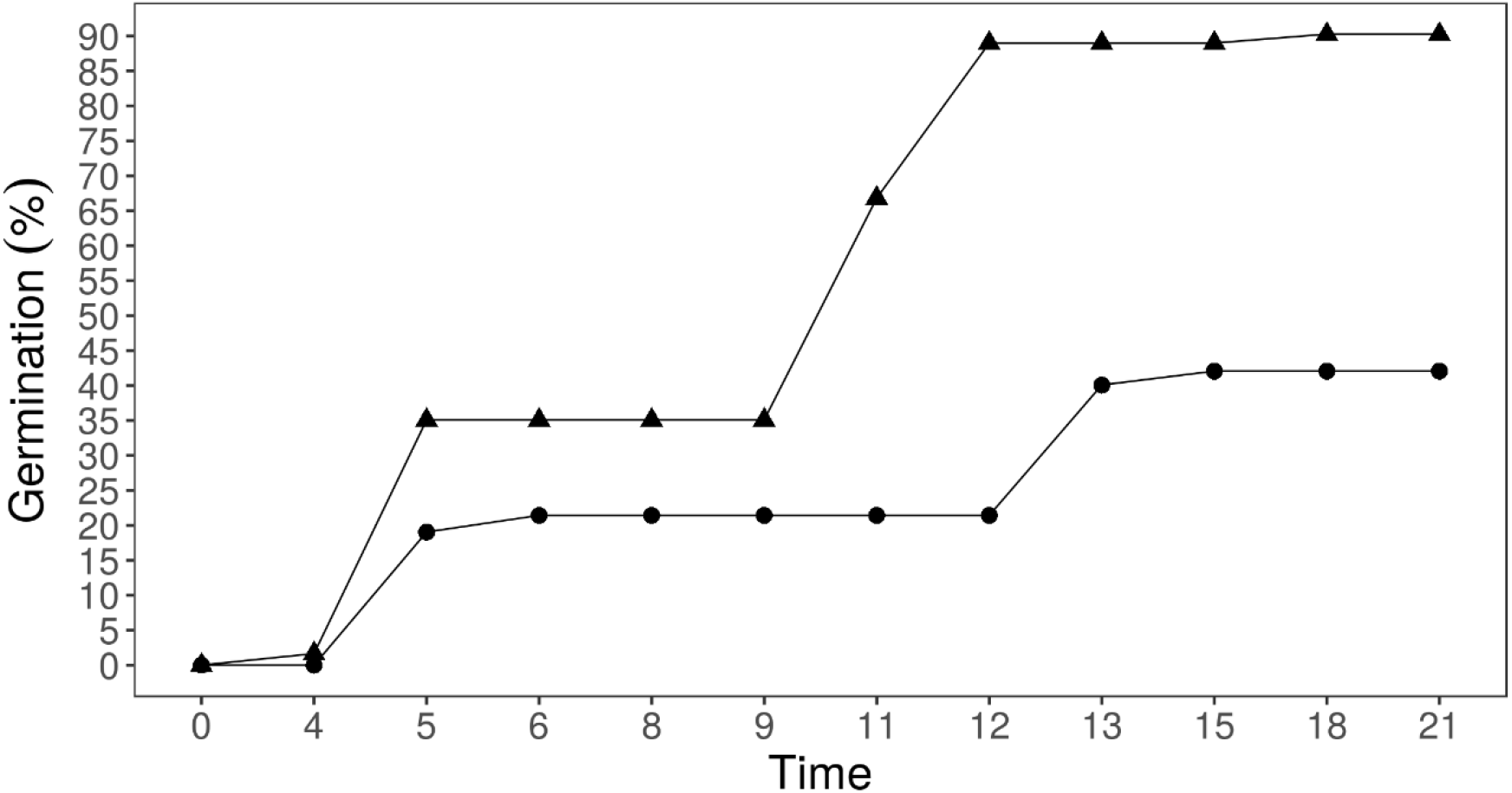
Cumulative germination of *S. tubulosa* according to the substrate: vermiculite (▲), and organic-based (●).

In the literature, the germination rate found to this species comprehends the range of 40 and 60% depending on the adopted temperature when scarified seeds are sowed in vermiculite [11]. At the end of the analyzed period, we found a mean of 90.26% germination, a greater value than the mentioned range of germination. Sales [11] measured germination 8 days after sowing, if we analyze this author provided range with the germination obtained on the eighth day (Figure 6), we obtained a corresponding value of this mentioned range. Another important factor is the environmental conditions that could influence seed germination. This author used continuous light and a constant temperature. These conditions could also be responsible for the difference between the mentioned values and the study data.

## 4. CONCLUSION

The parent population could affect the seedling production process, but this influence does not substantially preclude seedling production. The mechanical scarification procedure seems not to enhance seed germination at any of the studied indexes during the analysis period. Vermiculite promoted a better germination rate when compared to the organic-based substrate.

## Acknowledgments

Secretaria do Verde e Meio Ambiente of São Paulo due to its trainee programs. Also, the workgroup of Harry Blossfeld Municipal plant nursery, especially Luis and Angelica Tinoco, due to their insight on the importance of this work execution.

